# Estimating Speed-Accuracy Trade-offs to Evaluate and Understand Closed-Loop Prosthesis Interfaces

**DOI:** 10.1101/2022.06.13.495789

**Authors:** Pranav Mamidanna, Jakob L Dideriksen, Strahinja Dosen

## Abstract

**Objective:** Closed-loop prosthesis interfaces, combining electromyography (EMG)-based control with non-invasive supplementary feedback, represent a promising direction to develop the next generation of user prosthesis interfaces. However, we still lack an understanding of how users make use of these interfaces, and how to evaluate competing interfaces. In this study we use the framework of speed accuracy trade-off functions (SAF) to understand, evaluate and compare the performance afforded by two closed-loop user-prosthesis interfaces.

**Approach:** Ten able-bodied participants and one amputee performed a force matching task in a functional box-and-blocks setup at 3 different speeds. All participants were subject to both interfaces in a crossover fashion with a one-week washout period. Importantly, both interfaces used (identical) direct proportional control but differed in the feedback provided to the participant – EMG feedback vs force feedback. We thereby estimated the SAFs afforded by the two interfaces, and additionally sought to understand how participants planned and executed the task in the various conditions.

**Main results:** We found that execution speed significantly influenced the performance, and that EMG feedback afforded better performance overall. Notably, we found that there was a difference in SAF between the two interfaces, with EMG feedback enabling participants to attain higher accuracies faster than Force feedback. Further, both interfaces enabled participants to develop flexible control policies, while EMG feedback also afforded participants to generate smoother more repeatable EMG commands.

**Significance:** Overall, the results indicate that closed-loop prosthesis interfaces afford subjects to exhibit a wide range of performance, which is affected both by the interface and the execution speed. Thereby, we argue that it is important to consider the speed accuracy trade-offs to rigorously evaluate and compare (closed-loop) user-prosthesis interfaces.

## Introduction

Myoelectric interfaces that leverage electromyographic (EMG) signals recorded non-invasively from the residual muscles of amputees enable control of advanced upper limb prosthetic devices. These interfaces have been combined with supplementary feedback using non-invasive vibrotactile or electrotactile stimulation and principles of sensory substitution, to provide users with useful information regarding the state of the prosthesis. Together, these approaches promise to address a key challenge of closing the user-prosthesis loop to create the next generation of non-invasive interfaces aimed at improving the reliability and intuitiveness of controlling prostheses [1], [2].

A key limitation in the development of such closed-loop interfaces is a lack of more basic understanding of the role of supplementary feedback in the user-prosthesis interaction [3]. Researchers in the field have used tools and concepts from human motor learning and control to better understand how subjects integrate supplementary feedback to plan and control their devices. Consequently, supplementary feedback has been shown to aid learning internal models of the prosthesis [4], [5], improve state estimation [6] and psychosocial aspects of subjective experience [7], [8]. This knowledge was successfully applied to design better interfaces and to evaluate existing interfaces [9]. Despite these promising recent developments, the understanding of motor control in the context of prosthesis use is still in its infancy.

In a recent study, we showed how subjects could take advantage of supplementary feedback to develop flexible prosthesis control policies and exhibit a speed-accuracy trade-off [10]. Speed-accuracy trade-off is a ubiquitous behavioral phenomenon, observed in several species and across several tasks from foraging to tool use [11]. The speed-accuracy trade-off function (SAF) has been used as an instrument to understand both perceptual and motor ability and has a wide reception in the field of human-machine interfaces, building on seminal work by Fitts [12]. A variety of tasks inspired by this experimental paradigm have been applied in the context of myoelectric control [13]–[20]. In classical Fitts’ style pointing tasks, participants are required to move a cursor to a target location, specified by a target width and distance, and their movement time is recorded. Thereby, these experiments determine speed (movement time) as a function of task difficulty, while accuracy in such tasks is a given and correspond to “asymptotic” performance. Alternatively, one could hold task difficulty constant, and measure how the accuracy changes when the same task is performed at different speeds, a framework that has been successfully used in understanding motor skill [21]–[23].

A SAF so measured can be characterized by its intercept, rate, and asymptote, without making any assumptions on the functional form of the trade-off, barring monotonicity [24] (see Figure 1). The intercept characterizes the minimum time required to have any chance of success, the rate provides information about how rapidly the trade-off between speed and accuracy can be achieved, and the asymptote characterizes a performance ceiling when one performs a task slowly and carefully. Therefore, SAF has been proposed as a preferred metric to measure and understand a participant’s overall performance and motor ability [22], [24]. However, no current user-prosthesis interface has been analyzed using this methodology.

**Figure 1:**
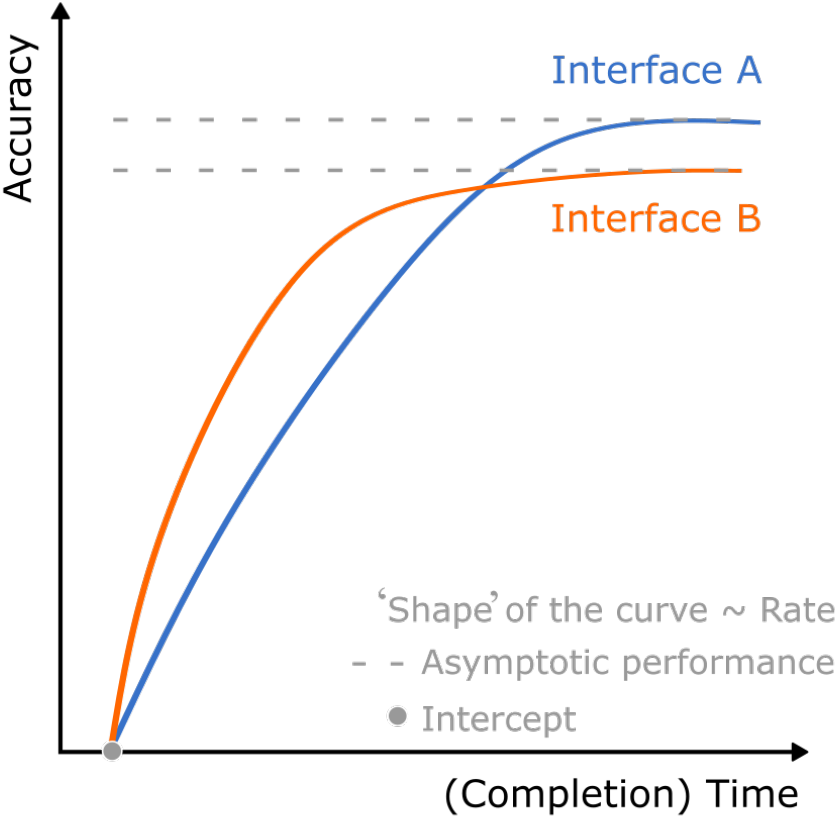
Speed-Accuracy Trade-off. A cartoon depicting the concept of speed-accuracy trade-offs as characterized by (1) intercept, (2) rate and (3) asymptotic performance, for two different interfaces.

A common practice in the field to evaluate (the effectiveness of) interfaces involves measuring performance in a given task at a ‘comfortable pace’. We argue that such an evaluation, which corresponds to sampling the SAF at a single point, is an insufficient indicator of the range of performance afforded by the (closed-loop) interfaces. Moreover, a comparison of competing interfaces is compromised when the comparison is based on a single point on the SAF. Such a comparison is limited in scope (a single point vs. a full SAF) and it could even entail comparing different points while assuming they are the same (a ‘comfortable pace’ might differ across subjects, tasks, and interfaces). Determining the SAF, on the other hand, allows a comprehensive characterization of performance and can provide unique insights that can be used to make informed choices. For example, consider two hypothetical interfaces shown in Figure 1. Sampling the two interfaces at different points of their respective SAFs leads to different conclusions about which interface affords better performance. Moreover, a user who emphasizes speed may be better off with interface A, but relaxing this requirement suggests interface B is a better bet, information which is only available through the SAF. Such a comprehensive assessment becomes even more pressing as there are several promising user-prosthesis interfaces that use different (combinations of) control (e.g., direct proportional, pattern recognition, regression etc. [25]) and feedback interfaces (e.g., force, aperture, proprioceptive feedback using different modalities [26]). Narrowing down the focus to closed-loop control of grasping force, arguably the critical function of hand prostheses, several (feedback) interfaces have been proposed in the literature [3], [26], [27]. However, comparisons of these interfaces are difficult since the performance is sampled at a single, and possibly different, point along the SAF.

In this experiment, we empirically study the SAF in closed-loop myoelectric control, using the prosthesis force-matching paradigm in a functional task – the box and blocks test – to (1) show how SAF can be used to evaluate (closed-loop) prosthesis interfaces and (2) thereby understand how they affect users’ ability to control the prosthesis. Specifically, we compare two interfaces which both use direct proportional control to modulate prosthesis velocity but differ in the feedback they provide to the subject – EMG feedback [28]–[30] vs force feedback (see Table A1 in [3]). We use a prosthesis force-matching paradigm to understand how well the two interfaces enable participants to achieve the same target force at three different speeds, ordinally defined as fast, medium, and slow (see Methods: Experimental Design). Since the difficulty of the task itself and control interface are fixed, the performance differences that arise from this experiment are a consequence of the feedback interfaces. Having sampled the SAF at the three distinct speed requirements, we investigate how the SAF differ for the two interfaces and analyze how the participants’ control policies change both across interfaces and speeds. Finally, we investigated if the results extend to amputees, using a case study of a single amputee.

## Methods

### Participants

Ten healthy, able-bodied participants (7 male and 3 females with a mean age of 28 ± 2 years) and one transradial amputee (female, 49 years old, 10 years since traumatic amputation of non-dominant hand, limited daily use of a single DoF myoelectric prosthesis) were recruited. All participants signed an informed consent form before the start of the experiment. The experimental protocol was approved by the Research Ethics Committee of the Nordjylland Region (approval number N-20190036).

### Experimental Setup

The experimental setup is shown in Figure 2A. Able-bodied participants donned an orthotic wrist immobilization splint, to produce near-isometric wrist flexion and extension, and the prosthetic device (Michelangelo hand, OttoBock, DE) was attached to the splint, with the arm placed in a neutral position. A custom-fit socket was made for the amputee. Two dry EMG electrodes with embedded amplifiers (13E200, Otto Bock, DE) were placed over the wrist flexors and extensors of the right forearm, located by palpating, and visually observing muscle contractions. Five vibrotactors (C-2, Engineering Acoustics Inc.) were positioned equidistantly around a cross-section of the upper arm and an elastic band was used to keep them in place. A standard Box and Blocks setup was used for the experimental task. The task instructions were shown on a computer screen placed at a comfortable viewing angle and distance. The prosthesis was connected to a standard laptop PC through a Bluetooth link, while the vibrotactors were connected through a USB port. The control loop for the experiment was implemented in MATLAB Simulink using a toolbox for testing human-in-the-loop control systems [31] and operated on the host PC in real time at 100Hz through the Simulink Desktop Real Time toolbox.

**Figure 2:**
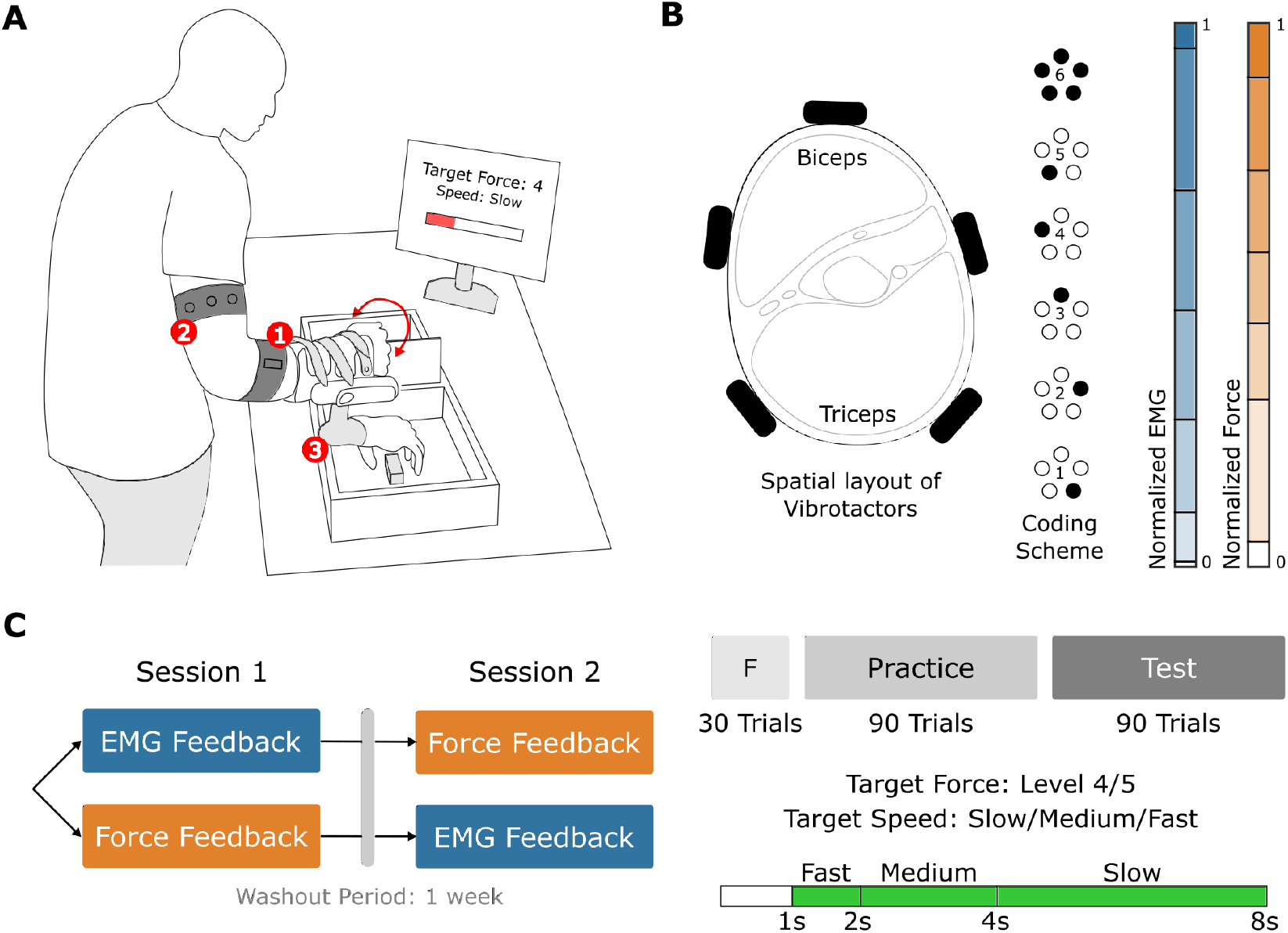
Experimental setup and protocol. (A) Sketch of the experimental setup showing 1. Two dry EMG electrodes placed on the forearm, 2. Vibrotactor array for delivering feedback placed on the upper arm and 3. The Michelangelo prothesis. (B) Vibrotactor array arrangement and coding scheme used for the feedback interfaces. Bars indicate how the normalized EMG and Force range was discretized to provide feedback. (C) Experimental protocol indicating the design (AB-BA crossover), trial structure and force and speed targets.

### EMG Control

Participants used near-isometric wrist flexion and proportional control to generate velocity commands to close the prosthesis. Opening the prosthesis was triggered by a strong contraction (see below) of the wrist extensors, instead of proportional control, since fine control of the opening was not relevant for the study. Two electrodes, placed on the flexors and extensors as explained above, were used to record the root mean square of the windowed (100 ms) EMG signal at 100Hz through the embedded prosthesis controller. The signals were subsequently filtered digitally using a second order Butterworth low-pass filter with a 0.5 Hz cutoff. The EMG envelope from each of the electrodes was normalized to 50% of the maximum voluntary contraction (MVC). For the flexor EMG, this corresponded to the maximum closing velocity of the prosthesis. A piecewise linear mapping between EMG amplitude and closing velocity was used to design the proportional controller, to compensate for the higher variability in the EMG signal at higher amplitudes (stronger contractions). The breakpoints for the mapping were defined as follows: EMG = {0.01, 0.1, 0.27, 0.47, 0.69, 0.95, 1}, velocity = {0, 0.25, 0.42, 0.59, 0.76, 0.9, 1}. For the extensor however, participants simply needed to reach 0.4 on the normalized range (corresponding to 20% MVC) to trigger hand opening.

### Vibrotactile Feedback Interfaces

In this study, we compared two feedback interfaces – EMG feedback and Force feedback. Both interfaces were identical in terms of the hardware and encoding (described below) and differed only in the variable which was provided as feedback – participants’ own EMG command vs prosthesis force. Five vibrotactors were placed circumferentially and equidistantly on the upper arm around a cross section containing the biceps. An elastic band was used to keep the tactors in place. A spatial encoding scheme consisting of six discrete levels of the feedback variable (EMG command or grasping force) was used for both interfaces. The first five levels were indicated by activating one of the tactors from the array while the sixth level was conveyed by activating all the tactors simultaneously (Figure 2B). If the vibrotactors evoked an unpleasant or poorly localized sensation, their position was adjusted until the participants could easily distinguish all six stimulation patterns (levels). The vibration frequency for all tactors was set to 200 Hz, and the stimulation pattern was updated at 50 Hz.

### EMG Feedback

In this interface, the participants were provided feedback about the EMG signal that they generated using their flexor muscles to control the closing velocity of the prosthesis. The six discrete levels were defined using the breakpoints of the piece-wise linear mapping described in section “Methods: EMG Control.” Therefore, as soon as the participants started contracting their wrist flexors, they received feedback about the EMG level (1-6) they were generating, thereby enabling them to predictively modulate to the target level. The breakpoints of the piecewise mapping were designed in such a way that if the participants reached a particular level of EMG (with the object contact established and stable), they will have applied the same level of force on the object. For instance, if a participant would generate and maintain EMG level 2, the prosthesis would close around the object and exert the level 2 of the grasping force (level boundaries defined in the next section).

### Force Feedback

The force applied on the blocks was measured by a sensor embedded within the prosthesis. The measured force, sampled at 100Hz by the embedded controller, was normalized and divided into six discrete ranges (levels) with boundaries at {0.05, 0.31, 0.45, 0.58, 0.73, 0.9 and 1} on the normalized scale. With this feedback interface, the participants received feedback on the level of force (1-6) applied on the object. Contrary to EMG feedback, where vibrotactile stimulation was delivered as soon as the myoelectric signal crossed the threshold of the dead-zone (e.g., when the prosthesis started closing), in the case of force feedback, the stimulation was delivered only after contact was established with the object.

### Experimental Design

The experiment was designed as an AB-BA crossover trial over two sessions with a one-week washout period between the sessions (Figure 2C). Half of the participants started with EMG feedback interface in Session 1 and switched to Force feedback in Session 2, while the other half of the participants did the opposite. A crossover design has been selected to control inter-group variability. In each session, the participants were instructed to perform the box and blocks test with two additional constraints, i.e., in each trial, they were required to (1) apply a specified level of force on the object (two levels of force were chosen as target forces – levels 4 and 5, see [10]), and (2) reach the target force within a specific time window. Thereby, we determined the speed-accuracy trade-off in a prosthesis force-matching task.

To adequately sample the SAF, participants were required to perform the task in three speed conditions – slow, medium, and fast, where each condition specified the time window for task completion. During the Slow condition, trials had to be completed within 4 – 8s, while for the Medium and Fast conditions the speed/time requirements were 2 – 4s and 1 – 2s, respectively. The time windows have been defined to capture the relevant domains of the SAF curve. Previous studies suggest that participants in a fast routine grasping task spend around 2s to achieve required force while they attain close to 100% accuracy at around the 6s mark [10]. In effect, we used a time-band methodology to derive the SAF [24]. While there exist several methodologies to obtain the SAF [11], [24], we believe that this approach reduced inter-subject variability in learning feedback control. This would not have been the case in, e.g., a deadlines-based methodology, where participants may have no incentive to perform the task at a slower speed if they were satisfied with their accuracy while using faster speeds.

The amputee subject followed the same protocol as the able-bodied participants, starting with Force feedback in Session 1 but returned 3 weeks later (as opposed to one week) to perform the task with EMG feedback.

### Experimental Protocol

Initially, all equipment (EMG electrodes, vibrotactors, wrist immobilization splint and the prosthesis) were placed on the participant. Then a brief calibration and familiarization followed in both sessions. During the EMG calibration phase, three 5-second-long maximum voluntary contractions (MVC) for both the flexors and extensors were recorded to calibrate the control interface. The MVC measurements were recorded in the same posture that the participants would use to perform the box and blocks task (similar to [32]), to address the effect of arm posture and prosthesis weight on the recorded EMG. Next, the participants were familiarized with the interface and were guided to explore how their flexor EMG signal affected the prosthesis closing velocity and how their extensor EMG signal triggered hand opening. Finally, they were familiarized with the vibrotactile feedback (common across both feedback interfaces) by performing a spatial discrimination task where they were presented with two sets of 18 stimulation patterns (3 repetitions x 6 levels, Figure 2B) and asked to identify the patterns. The experiment proceeded after ensuring that the participants achieved at least 95% success in the discrimination task, which normally took less than 5 mins.

After familiarization with the control and feedback, the participants performed 30 trials (10 per speed condition) of the modified box and blocks test to practice the time-constrained force-matching task. Each trial began by displaying the force and speed targets. The participants then had to modulate their muscle contraction and use the feedback interface to successfully complete the trial. Once the participant felt they successfully reached (or overshot) the target, they were instructed to extend their wrist to trigger hand opening. Immediately after the trial ended, the participants received knowledge of performance, which indicated if they achieved, overshot, or undershot the target force and target speed. During the practice trials, the participants were explained how to modulate their muscle contraction to control the closing velocity of the prosthesis. They were also clearly instructed to avoid eccentric behavior, e.g., in the slow condition they were instructed against holding their contraction at a low level until 4 s and then quickly correcting upwards, hence inadvertently making a fast/medium condition trial.

After the initial practice trials, the participants performed 90 training and 90 test trials with a break after every 30 trials. In each such block of 30 trials, the target speeds (slow, medium, and fast) remained the same for 10 trials, while the target forces (4, 5) were presented 5 times each in a random order. In addition, during the first 60 training trials, the speed targets were presented in a specific order – slow, medium, and fast – while during the remaining trials, this was also randomized.

### Outcome Measures

During each trial, the EMG commands and force measurements were recorded and processed to obtain the primary outcome measures – reach time and trial success. from when the participant started generating the EMG input (above the dead-zone) to the time point where the maximum force was reached during the trial. A successful trial was therefore one where the reach time satisfied the speed requirement (1 – 2s for fast, 2 – 4s for medium and 4 – 8s for the slow speed), and the reached force was within the corresponding force interval (target level). The trials were aggregated per speed condition to obtain percent success rates (S). Subsequently, we computed the rate of trade-off in success rate (ΔS per second) during fast-to-medium and medium-to-slow conditions to evaluate how quickly the participants traded speed for accuracy. For each participant, we computed the rate of trade-off as the difference in success rates (ΔS) between successive speed conditions divided by the difference in the corresponding reach times. For example, the rate of trade-off for participant *p* for fast-to-medium transition was computed as (*S*_*p*|*med*_– *S*_*p*|*fast*_)/(*T*_*p*|*med*_– *T*_*p*|*fast*_), where *S*_*p*|*cond*_ is success rate and *T*_*p*|*cond*_ is the average reach time in the condition *cond*.

Further, to understand how the participants planned and executed the task in the different speed and feedback conditions we computed three behavioral metrics. Firstly, we calculated the number of force corrections in each trial that the participants made, by counting the number of distinct plateaus (longer than 250ms) in the force trajectory [10]. For example, during the slow condition with EMG feedback the amputee subject made 4 force corrections in the trial shown in Figure 3. Then, we analyzed the generated EMG commands, to understand if one feedback type could enable the participants to generate (1) smoother and (2) more repeatable EMG commands. To evaluate smoothness, we calculated the integrated squared jerk of each trajectory, normalized to the reach time. To measure the repeatability, we computed the trial-by-trial variability of the generated EMG commands. We first normalized all EMG trajectories to 200 time points between the start of the trial and reach time, then measured the standard deviation at each of the 200 time points. As the final measure of variability, we computed the median of the standard deviations across the time points since the distribution of the standard deviations was found to be often skewed.

**Figure 3:**
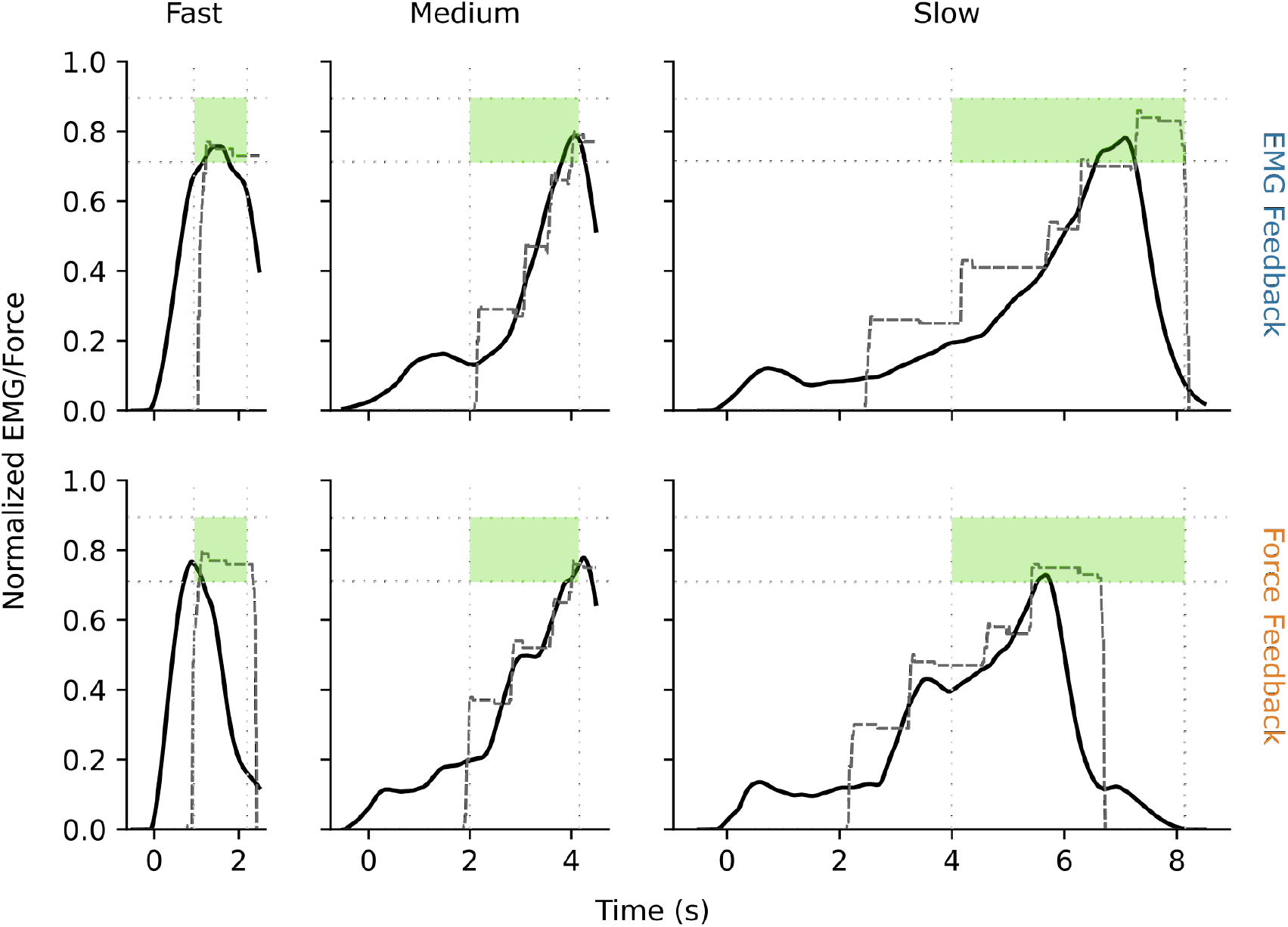
Representative Trials. Six representative trials (EMG commands in solid black, prosthesis force in dark gray) as performed by the amputee subject using the two different interfaces, at the three required speeds for target force Level 5. Faint dotted vertical and horizontal lines indicate time restrictions and force target bounds respectively. Green area depicts how trial success is determined as a combination of reaching the target force during the required time (speed).

### Statistical Analysis

Statistical analysis was performed on outcomes obtained during the 90 test trials. 3-factor mixed ANOVAs were fitted each for success rate, rate of trade-off and the behavioral metrics as the outcome, with two within-subjects factors – feedback interface and speed condition – and one between-subjects factor “order”, which denotes the order in which the participants were exposed to the feedback interfaces. We interpreted the main effect of order as an interaction between feedback interface and session, while the interaction effect between order and feedback interface was interpreted as the main effect of session, as is common in cross-over designs [33]. The assumptions of Normality, homogeneity of variance and sphericity were verified using Shapiro-Wilk’s, Levene’s and Mauchly’s tests, respectively.

Post-hoc analyses for differences in success rates between the two feedback interfaces at a given speed condition and between speeds for a given feedback interface were performed by using pairwise t-tests, adjusted using the Holm-Bonferroni method. The threshold for statistical significance was set at p < 0.05. Mean ± standard deviation of outcomes per group of interest are reported throughout the paper, unless noted otherwise.

## Results

### Representative Trials

Figure 3 shows representative trials of the amputee subject in all target speeds, with level 5 as the force target. Firstly, we can notice that both feedback types allowed the subject to flexibly control the prosthesis at different speeds and still succeed in the task, i.e., reaching the target grasping force within the given time window. Then, we can notice that the subject was slightly faster when using Force feedback than EMG feedback (especially noticeable in the slow trials), a feature that also holds across participants (see Figure 4A). Secondly, we can observe a difference in the “quality” of the generated EMG commands across the feedback conditions. While the EMG commands produced during the fast condition is largely similar between the feedback types, the EMG signal generated during both medium and slow trials is smoother during EMG feedback as opposed to Force feedback, where the EMG commands exhibit distinct “jumps”.

**Figure 4:**
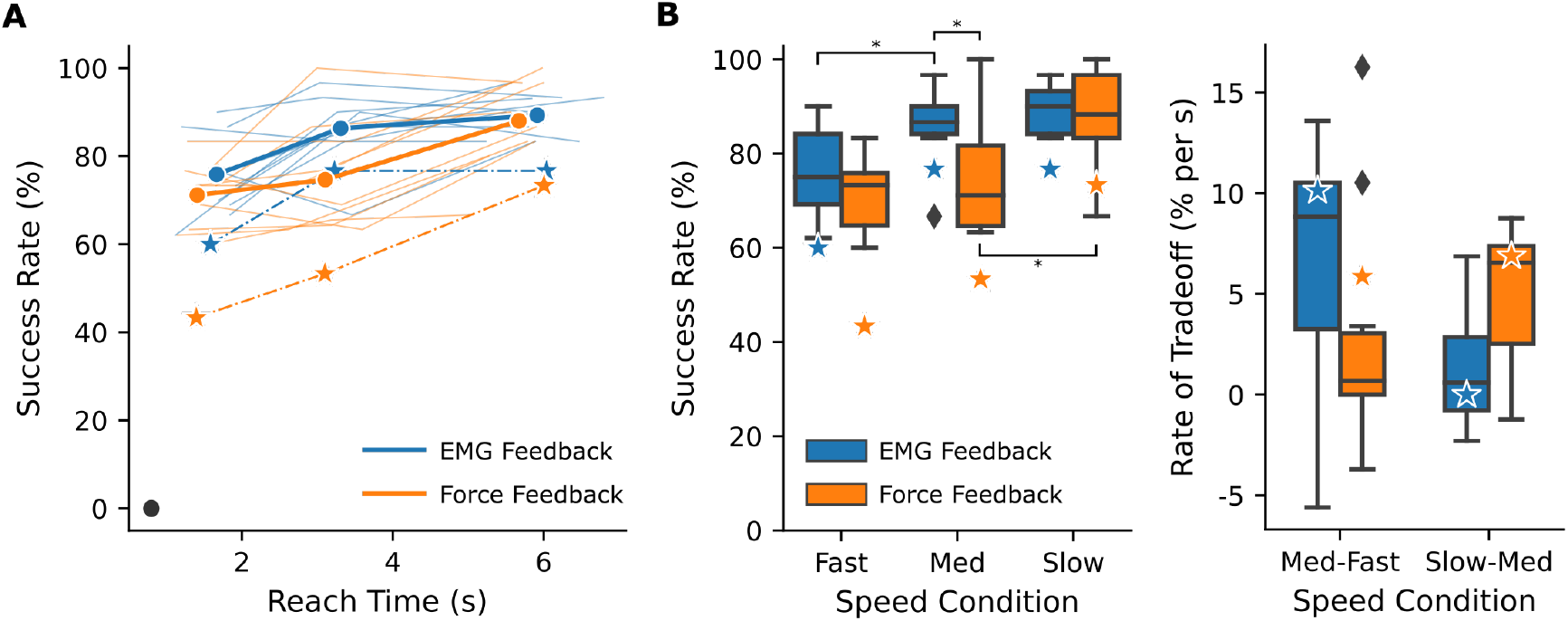
Speed-Accuracy trade-offs in prosthesis force control. (A) Individual speed-accuracy trade-off curves are plotted for each participant (faint lines), group means (bold lines) and the amputee subject (dashed lines and stars). Bold diamond indicates the time (X-intercept) when success is zero. (B) Same as A, but box plots show success rates of all participants during each of the ordinal target speeds (left). Box plots showing rate of trade-off (% per s) across the target speed transitions. Colored stars represent results of the amputee subject.

### Speed-Accuracy Trade-offs

The participants’ speed-accuracy trade-off curves showed a general tendency to be monotonic (14 out of 22, Figure 4A), and when they were not monotonic, they were only off due to a few trials (1 – 3 trials), while the mean SAF across participants were monotonic. This was true for both feedback interfaces. Next, we fit a 3-factor ANOVA by treating the speed condition as categorical to analyze the effect of feedback interface and speed condition on success rate. We observed a significant effect of feedback interface (p=0.006), and speed condition (p=2.3×10^−6^) on success rate as well as a significant effect of session (feedback interface x feedback order interaction effect, p=0.003).

We then analyzed if feedback interface affected performance at each of the speed conditions. In the Fast condition, we did not observe a significant effect of feedback interface (EMG: 75.8 ± 9.4%, Force: 71.1 ± 7.4% see Figure 4B), while in the Medium condition, we observed that participants performed significantly better using EMG feedback than Force feedback (EMG: 86.3 ± 8%, Force: 74.6 ± 12.2%, p-adj=0.022). In the Slow condition (asymptotic performance), as expected, we observed that the interface had no significant effect on performance (EMG: 89.2 ± 4.8%, Force 88 ± 10.2%). Taken together, we see that while the feedback interface had a significant effect on success rate overall, it was in the Medium speed condition that this difference originated from. Further, EMG feedback enabled participants to reach asymptotic performance sooner, with participants significantly improving their performance between Fast and Medium conditions (p-adj=0.03) but not between Medium and Slow conditions. On the contrary, participants exhibited significant improvement between Medium and Slow conditions (p-adj=0.004) while using Force feedback. The two feedback types are therefore characterized by SAFs that are qualitatively different, while still allowing similar asymptotic performance.

Therefore, we analyzed if the observed rate of trade-off in success rate (% per s) for the Fast to Medium, and Medium to Slow transitions were significantly different between EMG and Force feedback (see Figure 4B, right). Visually, there appears to be a difference in both cases, with the higher rate of trade-off for EMG feedback during Fast to Medium (6.6 ± 6.3% vs 2.7 ± 6.1% per second) and opposite for Medium to Slow transition (1.3 ± 2.8% vs 5.1 ± 3.5% per second). However, the difference was not statistically significant.

Performance of the amputee subject followed the trends of the able-bodied participants (Figure 4, stars). While the asymptotic performance was nearly identical (EMG: 76.6%, Force: 73.3%), the amputee participant reached higher success rates with EMG feedback in both Fast and Medium conditions, with the largest difference in the latter (Medium condition: EMG: 76.6%, Force: 53.3%).

### Behavioral Analyses

We sought to understand the behavioral differences between the feedback types, i.e., how the different interfaces allowed participants to plan and execute movements (Figure 5). First, we investigated if participants developed different strategies during the different speed targets. We found that both feedback interface (p=0.005) and speed condition (p<1e-15) had a significant effect on the number of corrections made by the participants (Figure 5A), and the feedback and session exhibited significant interaction (p=0.02). Therefore, the participants were able to flexibly modify their control policies by using the available feedback, especially during the Medium (2.2 ± 0.3, 1.8 ± 0.5 corrections per trial) and Slow conditions (3 ± 0.4, 2.8 ± 0.3 corr. p/trial) compared to the Fast condition (0.5 ± 0.3, 0.2 ± 0.2 corr. p/trial).

**Figure 5:**
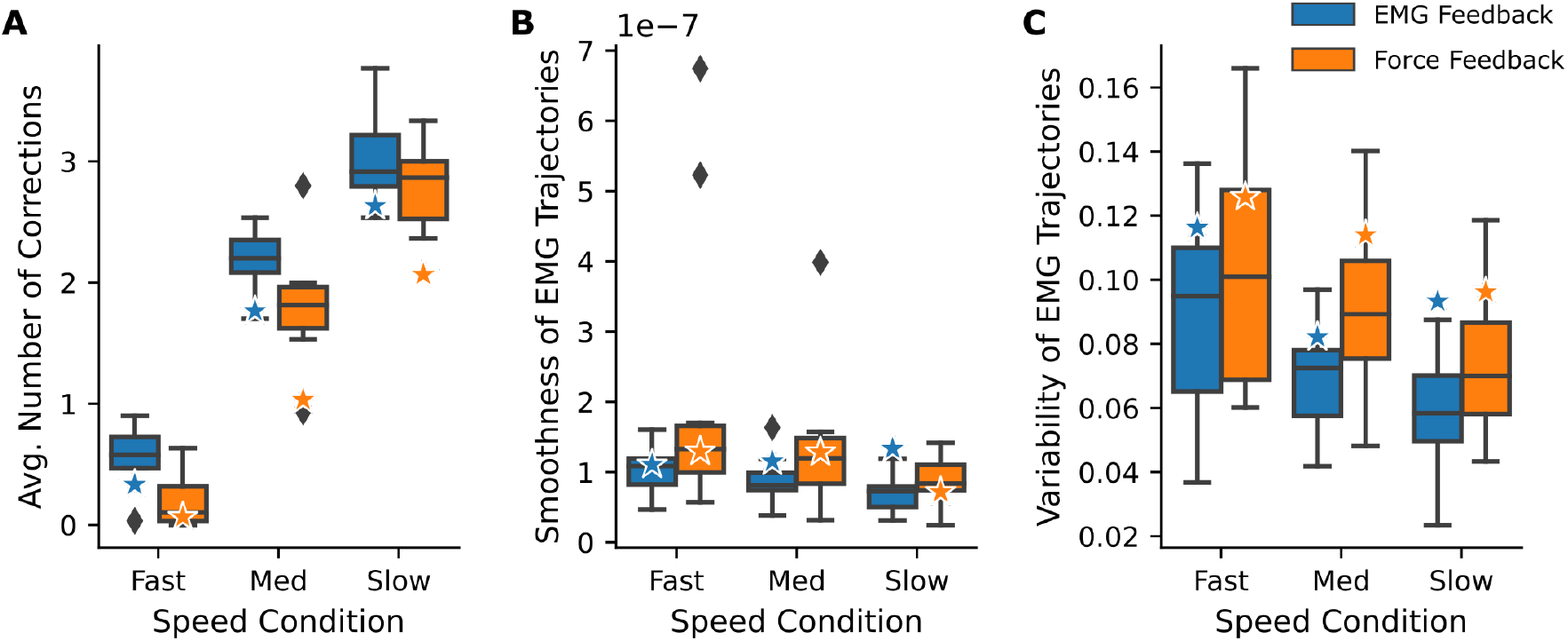
Behavioral metrics for both interfaces, across participants. (A) Average number of force corrections (distinct force plateaus) per trial. (B) Smoothness of EMG trajectories (commands) generated by the participants, computed as integrated squared jerk of the normalized EMG amplitude. (C) Trial-by-trial variability of EMG commands generated by the participants. Stars represent results of the amputee subject.

Next, we analyzed the generated EMG commands by measuring the smoothness and trial-by-trial variability (Figure 5B, C). We found that the feedback interface had a significant effect on both metrics (p=0.03 for smoothness, p=0.002 for trial-by-trial variability). That is, EMG feedback enabled the participants to make smoother and more repeatable commands compared to Force feedback. Additionally, the speed condition had a significant effect on both metrics (p=0.01 for smoothness, p=0.0006 for variability), while the session significantly influenced trial-by-trial variability (p=0.001).

The behavior of the amputee subject followed the results of able-bodied participants. However, the smoothness of EMG commands with EMG feedback was worse than with Force feedback in the Slow condition.

## Discussion

Speed and accuracy are critical factors in the context of human-machine interfaces. Investigating speed-accuracy trade-off functions provides a thorough understanding of task performance and motor ability but has not been applied to study user-prosthesis interfaces. Here, we empirically derived the SAF using a prosthesis force-matching paradigm in a functional box-and-blocks task for two different closed-loop interfaces, which only differed in the feedback provided to the participants – EMG feedback vs Force feedback. Expectedly, the speed at which participants performed the force-matching task imposed a trade-off with accuracy regardless of the feedback type. However, the SAF was different for the two interfaces, as EMG feedback substantially outperformed Force feedback in the Medium speed condition and thereby enabled participants to reach asymptotic performance sooner. In addition, we found that EMG feedback enabled smoother and more repeatable EMG commands. Therefore, the results demonstrate that the SAF methodology can provide crucial insights regarding both evaluating and understanding of closed-loop interfaces for prostheses control even in functionally relevant task settings.

### SAF to Evaluate Closed-Loop Interfaces

Evaluation and comparison of user-prosthesis interfaces is challenging and multi-faceted. Despite rapid development of promising control and feedback interfaces [25], [26], their comparison has received less attention, barring a few exceptions [29], [34]. While it is a difficult undertaking due to various reasons such as incomparable experimental setups and tasks, here we showed that it is additionally compounded by measuring the performance only at a single speed (sampling at a single point on the SAF). For example, if we had only measured performance in the Fast condition in this study, we would infer that both interfaces enable similar performance, while they are in fact significantly different when used at the Medium speed. We argue therefore, that it is valuable to compare interfaces at more than a single point on the SAF especially since the shape of the SAF afforded by different interfaces is unknown.

Here, we used the SAF framework to rigorously compare two closed-loop interfaces in a functional force-matching task. By enforcing task execution at different speeds, we elicited a range of success rates that were significantly affected by the feedback interface used. We expected that EMG feedback would enable better success rates during the Fast condition, since it promotes predictive control [28], [29], but that was not the case. We believe that this is likely due to two reasons. First, Fast condition might have been too restrictive, with a short 2 s window, for the participants to exploit the EMG feedback effectively for online adjustment of control commands. Second, the task included only 2 force levels and the participants received training before performing the test trials. The training might have enabled participants to acquire a reliable internal model and achieve a good performance when using Force feedback despite the short time window (which basically precluded the use of force feedback to drive the corrections). However, we noticed a large difference in success rates between the two interfaces in the Medium condition. Therefore, the results demonstrate that the expected advantage of EMG feedback over Force feedback occurs in this range of movement speeds, where the former allows users to predictively modulate their contractions to reach the target level as opposed to ‘reactively’ jump between levels. Finally, the feedback interfaces resulted in similarly high performance in the Slow condition, as the participants had enough time to reach the goal by focusing on either of the two feedback signals. The present study therefore demonstrates that SAF allows identifying the time interval in which feedback (Force or EMG) becomes an important factor for the effectiveness of the control loop.

Taken together, we found that the asymptotic performance for both interfaces was similar, while EMG feedback allowed participants to approach the asymptotic performance sooner. Note that this important characterization of the two feedback types is derived from the trade-off itself and could not be obtained if the performance was assessed in a single point. More generally, SAF provides a way to estimate the expected completion time to guarantee a given (e.g., 90%) performance in a task, and therefore can be a relevant instrument for meta-analytic comparison of interfaces across studies. Moreover, we believe that determining SAF will be advantageous for person-based approaches to designing prosthesis interfaces [35], by e.g., determining the appropriate user-prosthesis interface for the amputee based on their inherent speed preferences (see Figure 1). Thereby, in the present study, we provided a holistic comparison of the performance afforded by two established interfaces in a functional task and added to a pool of methods that have been recently developed to assess the performance as well as behavior of the users of closed loop prostheses [15], [36].

### SAF to Understand Closed-Loop Interfaces

Closed-loop user-prosthesis interfaces are a promising technology likely to translate into clinical applications, but currently still facing several conceptual and implementational barriers [1], [27], [37]. A key prerequisite for designing better closed-loop interfaces is to understand the complex interplay between feedforward and feedback control processes of the users, and how different interfaces facilitate it [3], [6], [10]. We believe that studying the SAF, as described here, is an effective instrument to approach this point as it enables us to understand how users interact and exploit different interfaces to achieve specific (time bound) goals.

Here, in addition to measuring performance, we used SAF to understand how participants planned and executed movements in a functional prosthesis task. We found that both closed-loop interfaces enabled users to develop flexible control policies. That is, participants were able to incorporate feedback to varying extents to guide their behavior during the different speed conditions, as reflected in the number of force corrections they made. Then, we found that EMG commands generated when participants used EMG feedback were smoother than when they used Force feedback. Interestingly, this suggests that even though participants received discretized feedback, they could exploit EMG feedback to predictively guide their contractions to generate an overall smooth control input. Combined with the low trial-by-trial variability across speed conditions, EMG feedback effectively reduced the uncertainty in generating prosthesis commands, a central aim of implementing supplementary feedback [6], [35]. Our results therefore add to a body of evidence which underscores the promise of some form of predictive feedback, about users’ own intention [5], [28], [29].

Together, the flexibility, smoothness and repeatability measures which are hallmarks of skilled behavior, help us understand how participants incorporate supplementary feedback in their control policies. And investigating the SAF provides a suitable framework for such an analysis. Finally, we also found that all outcome measures had similar trends in the amputee experiment. We believe that this is an encouraging result, albeit expected since motor planning and execution should remain comparable across able-bodied participants and amputees, especially when using a simple command interface (direct proportional control).

### Limitations and Outlook

A limitation of the current study is that we always required participants to make ‘strong’ contractions (30-45% MVC) to reach the target forces. However, the trade-offs (SAF) may be influenced by the force users want to generate. Another limitation is that while we performed a single session study to establish the conceptual framework of SAF, the shape of the SAF may change across days, in which case the SAF for both interfaces may become identical after practice, but this remains to be investigated.

Measuring the SAF can be an instrument for assessment of prosthesis control with rather general applicability. Future studies should be therefore conducted to investigate how the control interface (direct control, pattern recognition etc.) of the user-prosthesis loop affects the SAF, relative to the effects of feedback interface as explored here. In addition, this approach could be used to compare feedback interfaces which differ only in their encoding schemes (e.g., discrete vs continuous), while the feedback variable remains the same. The intercept (see Figure 1), which characterizes the minimum time required to have any chance of success, did not play a role in our current setup since the control interface was always the same (direct proportional control). However, in case one wishes to compare interfaces that allow different maximum velocities (e.g., due to different sensitivities for the proportional controller), or when one is required to change grips by co-contractions, it becomes crucial to understand the intercept as well. Finally, this framework can be extended to multi-dimensional task spaces, for example to characterize the trade-offs in prehension (posture matching with prostheses) combined with force-matching, to create better interfaces for current state-of-the-art commercial prostheses.

## Conclusion

In this study, we empirically derived SAF in prosthesis force control using a functional box-and-blocks task. We demonstrated that two closed-loop myoelectric interfaces which differed only in the variable provided as feedback to the participants – EMG feedback vs Force feedback – exhibited different SAFs. EMG feedback afforded better performance throughout, but especially at medium speeds, and enabled participants to develop stronger feedback control. We argue that the methodological advancement provided here is a valuable step forward in evaluating and understanding (closed-loop) user-prosthesis interfaces.

## Acknowledgements

We expressly thank Shima Gholinezhad for her help in organizing and conducting the experiments with the amputee subject. We thank Ana Sofia Santos Cardoso for their help in making Figure 1(A). This work has been supported by the projects 8022-00243A (ROBIN) and 8022-00226B funded by the Independent Research Fund Denmark.

## Notes

### Competing Interest Statement

The authors have declared no competing interest.

